# Unlocking Sensitive Data with SPHERE in the Age of AI

**DOI:** 10.64898/2026.09.01.748580

**Authors:** Zihuai He, Junyoung Park, Rafael Catoia Pulgrossi, Justin Lee, Robert R Butler, Audrey Weber, Lu Tian, Xiang Zhang, Julie Fangran Wang, Sharon Sha, Elizabeth C. Mormino, Tony Wyss-Coray, Victor W. Henderson, Frank M. Longo, James Zou, Manisha Desai, Russ Altman

## Abstract

Sensitive human data underpin discoveries across medicine, biology and the social sciences, yet privacy regulation often prevents sharing them with collaborators or artificial intelligence (AI) systems. We introduce SPHERE, a model-free method that makes sensitive datasets directly usable by AI and shareable for open science as a synthetic twin, while the original records never leave the local environment. Across 33 datasets spanning five scientific domains, SPHERE protects individual privacy against adversarial re-identification attacks while preserving the data’s statistical structure: means, variances and correlations are reproduced exactly, effect size and P value in linear statistical analysis are numerically identical, nonlinear machine-learning utility is retained, and each twin is generated in seconds on a laptop. Frontier AI agents running on the twin reach the same scientific conclusions as on the original records. Analyses of the twin reproduce genome- and proteome-wide results at UK Biobank scale and recover the findings of landmark studies across three independent cohorts and consortia. The approach also extends to deep-learning embeddings across language, vision and time-series, with minimal utility loss. We make the Stanford Alzheimer’s Disease Research Center cohort openly available for the first time, as a SPHERE twin spanning nine modalities that any registered researcher can analyze without an approval process. We release SPHERE with certification of each twin’s privacy and fidelity, and an AI agent that autonomously executes research tasks on sensitive data without ever accessing it. Sensitive datasets that are currently closed to research could thus become routine inputs to open science and AI to enable key discoveries.

---

Artificial intelligence (AI) has moved to the center of scientific discovery over the past decade. Foundation models trained on diverse, large-scale data have accelerated drug discovery^1^, enabled generalist clinical reasoning^2^, and begun to transform research across the physical and social sciences^3,4^. Model capability scales predictably with training data volume and diversity^5^, making access to rich, heterogeneous datasets a central constraint on what these models can achieve. AI agents, meanwhile, now conduct end-to-end research autonomously, designing experiments, writing and executing analysis code, and interpreting results, compressing weeks of expert work into hours ^6,7^. Yet these systems require access to the data being analyzed. Frontier models typically operate through cloud APIs that are not authorized to receive sensitive data under standard terms, while local open-source alternatives demand substantial infrastructure and sacrifice much of the capability that makes these agents useful. This bottleneck is most acute for the datasets that drive the most consequential questions, including electronic health records, national biobank measurements, clinical trial cohorts, government survey microdata, and financial transaction records. Such data cannot be freely handed to an AI system, used to train a model, or shared with collaborators without extensive governance, consent, or data-use agreements such as HIPAA^8^ or GDPR^9^.

This barrier has three practical implications. First, researchers holding sensitive records cannot send them to a cloud-based AI service under standard terms, foreclosing autonomous artificial intelligence systems. Second, multi-site collaborations need to pool individual-level data for adequately powered joint analyses, but direct transfer is prohibited even between trusted partners^10^. Federated learning partially addresses model training but cannot support arbitrary statistical analyses, requires substantial infrastructure at every site, and cannot produce a shareable dataset ^11^. Third, institutions holding data of broad scientific value must choose among restrictive access procedures, pre-specified summaries that cannot answer new questions, or full microdata release that risks re-identification^12,13^.

Synthetic data generation offers a path toward resolving all three, and many methods have been developed to this end^14^. Each, however, fails on at least one dimension critical for AI applications. Noise-based approaches (differential privacy, DP) add calibrated noise to achieve privacy guarantees^15^, but the noise dilutes weak associations, collapses statistical power, and degrades machine learning (ML) utility that downstream analyses depend on. Generative models (generative adversarial networks (GAN)^16^, diffusion models^17^) achieve competitive ML utility but do not preserve exact statistical inference, provide no formal privacy guarantee, and require dataset-specific training with substantial compute expense. Statistical sequential methods^18^ generate plausible records but do not pass empirical privacy tests and are sensitive to model misspecification. Exact statistical inference is structurally unavailable to this model-based class which fits a distribution to the data. No existing method simultaneously satisfies all five requirements that a drop-in replacement for sensitive data requires: principled privacy, exact distributional fidelity, exact statistical inference, competitive ML utility, and practical speed.

Here we present SPHERE (Fig. 1a), a synthetic twin dataset generator that satisfies all five requirements at once. SPHERE transforms the data rather than perturbing it, adding no noise, fitting no model and requiring no dataset-specific calibration. As a result, means, variances, and correlations are identical to the original by mathematical guarantee; linear statistical analysis, including every ordinary least squares (OLS) coefficient, standard error, *p*-value, and principal component analysis (PCA) loading reproduces the original to machine precision. Across 33 benchmark datasets spanning five scientific domains, SPHERE also retained competitive nonlinear ML utility, though not exact, while outperforming nine alternatives on the combined criteria of privacy, distributional fidelity, exact linear inference and practical speed. SPHERE runs in *O*(*np*) time, completing in seconds on a laptop, orders of magnitude faster than alternatives, and scaling to cohorts of hundreds of thousands of participants and millions of features that existing approaches could not previously reach.

**Figure 1.**
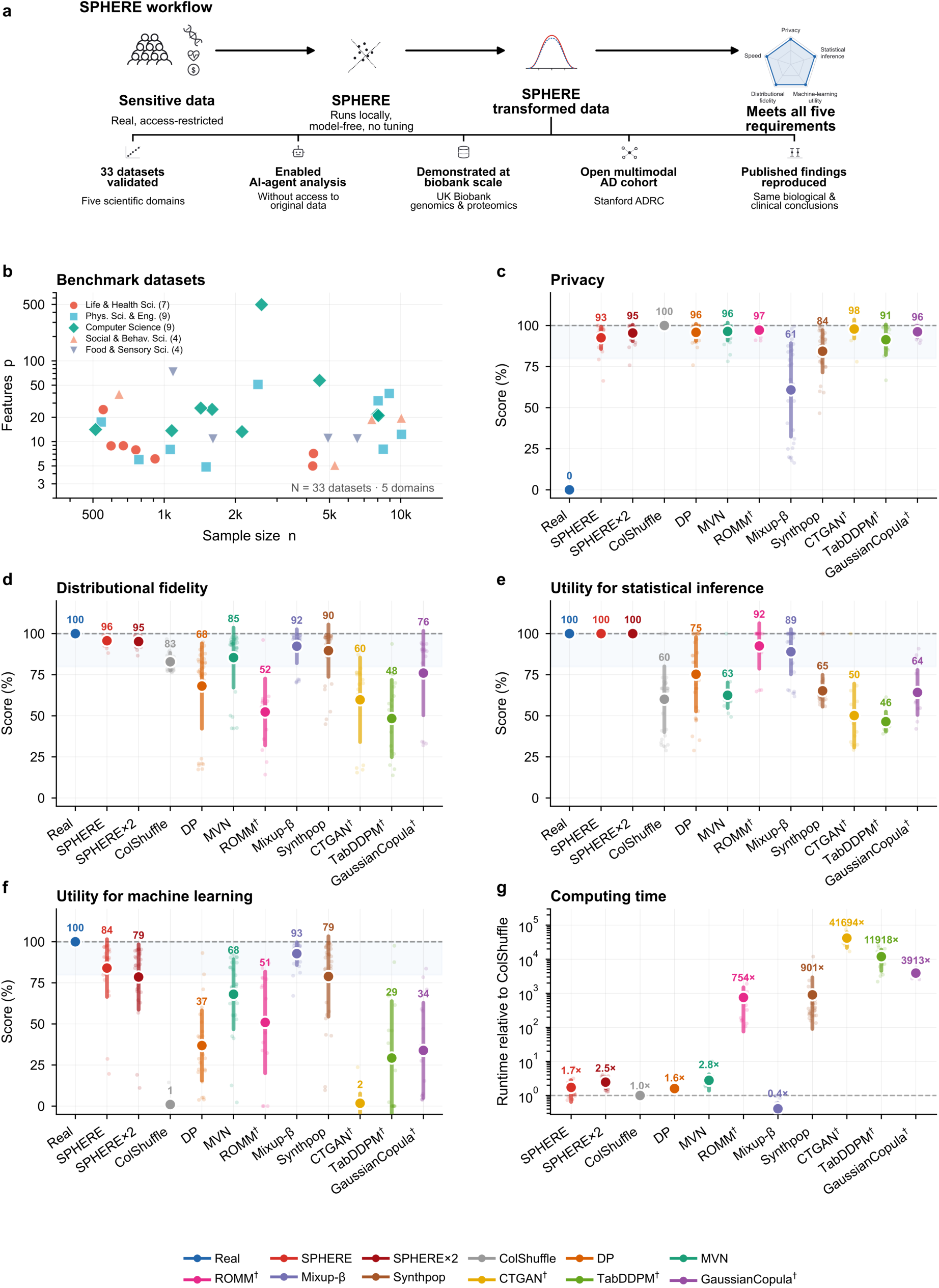
The SPHERE workflow and empirical evaluation of synthetic data methods on 33 real-world datasets spanning five scientific domains. **(a)** SPHERE workflow: real, access-restricted data are mapped to a twin dataset that is safe to share and publish while preserving exact linear statistics. The twin dataset is validated across 33 benchmark datasets, scales to population-scale biobank data, can be openly shared (Stanford ADRC), is usable by AI agents and deep-learning models, and reproduces published findings. **(b)** Benchmark datasets from OpenML plotted by sample size (*n*) and feature dimension *p* colored by domain. **(c)** Privacy composite from Anonymeter singling-out, linkability, and inference risk, normalized between real data (0%) and column shuffling (ColShuffle; 100%), with higher values indicating greater privacy. **(d)** Distributional fidelity from preservation of means, variances, correlations and Kolmogorov– Smirnov (KS) distributional agreement. **(e)** Utility for statistical inference (composite of ordinary least-squares (OLS) Type I error, power, confidence interval (CI) coverage). **(f)** Machine learning (ML) utility, calculated as a composite of Train on Synthetic, Test on Real (TSTR) *R*^2^ values across five models: Elastic Net (EN), Support Vector Machine (SVM), Random Forest (RF), Gradient Boosting (GB), Multi-Layer Perceptron (MLP) models. **(g)** Computing time relative to ColShuffle on a single Apple M3 CPU core, shown on a log scale. Dots show means across datasets, whiskers show ± 1s.d. and faded points show individual dataset scores. SPHERE×2 applies SPHERE twice in sequence for stronger privacy at the cost of reduced ML utility. MVN is a classical multivariate-normal parametric baseline. † ROMM, CTGAN, TabDDPM, and GaussianCopula evaluated on 16 datasets with *n* ≤ 2,000.

We use SPHERE in five applications that data-access barriers had previously blocked. First, frontier AI agents develop and execute analyses on sensitive data without accessing original records, with all AI-generated code that transfers unchanged to the real data (NHANES, *n*=4,899). Second, UK Biobank (UKB)-scale genomics (*n* = 461,307, 9,692,208 variants) and proteomics (n= 44,526; 2,916 proteins) discoveries are fully reproduced, showing that biobank-scale findings can be inspected, replicated and extended without exposing participant-level records. Third, we make the Stanford Alzheimer’s Disease Research Center (ADRC) cohort available for the first time under registered access, as a synthetic twin spanning nine modalities, paired with an agentic AI that allows users to browse, download, and run autonomous analyses directly. Fourth, published biomedical studies reproduce end-to-end from SPHERE data alone, yielding the same biological and clinical conclusions as analyses of the original records. Finally, SPHERE further generalizes to deep-learning embeddings across the language, vision, and time series architecture families, with minimal utility loss. We also deliver the SPHERE desktop application, a platform for generating, evaluating, certifying, and sharing SPHERE data with a sandboxed agentic AI for autonomous research, requiring no programming expertise.

## SPHERE simultaneously satisfies principled privacy, exact distributional fidelity, exact statistical inference, competitive ML utility, and practical speed

We evaluate SPHERE against nine comparison methods on 33 real-world datasets spanning five scientific domains (*n* = 522–10,000, *p* =5–490; Fig. 1b; Methods and Supplementary Table 1). No existing method simultaneously satisfies all five requirements (principled privacy, exact distributional fidelity, exact statistical inference, competitive ML utility, and practical speed), but SPHERE does (Fig. 1b-g). Every OLS coefficient, standard error, p-value, and PCA loading computed on SPHERE data is numerically identical to the value computed on the original data. Anonymeter privacy scores^19^ reach 92.5 ± 6.8% (two-anchor normalized, mean ± s.d. across 33 datasets; singling-out 99.2 ± 2.3%, linkability 91.2 ± 8.2%, inference 87.2 ± 17.6%) under SPHERE’s default settings, which were chosen to balance privacy against ML utility (Fig. S1; Fig. 1c). SPHERE preserves every mean, variance, and correlation exactly (*Δ* = 0 by construction), and attains the highest distributional-fidelity score of any method, where every other method shows dataset-specific distortion (Fig. 1d). SPHERE is the only method that simultaneously controls Type I error at the nominal level, preserves statistical power, and maintains correct confidence interval coverage (Fig. 1e). SPHERE matches real-data performance exactly for linear models (elastic net 100%) and retains 76-89% for non-linear models (random forest 76%, gradient boosting 84%, multi-layer perceptron (MLP) 83% and support vector machine 89%) (Fig. 1f). SPHERE× 2, which applies SPHERE twice in sequence (*k* = 2; SPHERE itself is a single application, *k* =1) offers stronger privacy (95.4 ± 5.0% versus 92.5 ± 6.8% for SPHERE) with identical distributional fidelity and exact statistical inference, at the cost of reduced ML utility (79% versus 84%) (Fig. 1c, f). SPHERE runs in *O*(*np*) time: 1.7× ColShuffle (SPHERE×2, 2.5×), versus 901× for Synthpop^18^, 11,918× for TabDDPM^17^, and 41,694× for CTGAN^16^ (Fig. 1g).

Every competing method underperforms on at least one of these five criteria, and on a different axis. DP-Gaussian^20^ achieves strong privacy but adds noise that collapses ML utility (37%). ColShuffle preserves marginal distributions but destroys all inter-variable associations, erasing downstream ML utility (1%) and lacks any formal guarantee. Random orthogonal matrix masking (ROMM)^21^ alters marginal distributions, failing distributional fidelity (52%), degrades per-sample feature structure resulting in poor ML utility (51%), and is computationally infeasible at AI-scale sample sizes (754×). Multivariate Normal (MVN) preserves means and variances under Gaussian assumptions but fails on non-Gaussian data (statistical inference 63%) and provides no formal privacy guarantee. Mixup-*β*^22^ preserves marginal fidelity (92%) yet offers the weakest empirical privacy among synthetic methods (61%). Synthpop^18^ and GaussianCopula^23^ generate plausible-looking records but lack a formal privacy guarantee and, respectively, fail exact inference (65%) and competitive ML utility (34%). CTGAN^16^ and TabDDPM^17^ require GPU-hours of dataset-specific training (current results reflect default training) and introduce systematic distributional distortion (60% and 48%) with no formal privacy guarantee (full per-dataset results in Supplementary Table 2-6; method details in Supplementary Note 2).

The properties of SPHERE come with trade-offs, which we state explicitly. Exactness applies to analyses determined by the first two moments; for models outside this class SPHERE empirically retains 76–89% of real-data performance in ML utility. Marginal distributions are preserved exactly for Gaussian and for categorically encoded columns, but SPHERE mapping smooths other continuous columns toward Gaussian; the Kolmogorov–Smirnov term is therefore the only component of the distributional-fidelity composite that falls below 100%, and the effect is on exploratory visualization rather than on inference. Privacy is established by reconstruction ambiguity together with empirical adversarial evaluation. Where stronger protection is warranted, applying SPHERE k times raises privacy at a measured cost in non-linear utility (95.4% versus 92.5% privacy, 79% versus 84% ML utility at k = 2), making the trade-off an explicit control rather than a fixed operating point.

## AI virtual labs reach consistent conclusions on SPHERE and original data

We next sought to demonstrate a key use of SPHERE, allowing AI agents to autonomously execute research tasks on sensitive data without ever accessing it. In Scenario I, a four-agent AI Virtual Lab (Principal Investigator, Epidemiologist, Biostatistician, Clinical Expert) was run on both SPHERE and original records under identical prompts, in US adults from NHANES 2017–2018 data^24^ (*n* = 4,899; 19 predictors; Fig. 2) to identify which risk factors independently predict systolic blood pressure. PCA confirms real and SPHERE point clouds are fully intermixed (Fig. 2a). Anonymeter privacy scores are near-ceiling on all three metrics (singling-out 100%, linkability 96%, inference 99%; Fig. 2b). Both synthetic and original datasets reached similar scientific conclusions and identified the same top predictors: age (*β* = +9.2 mmHg per 1-s.d. increase), Black race (+3.2 mmHg relative to the reference group), and BMI (+2.3 mmHg per 1-s.d. increase), with *R*^2^=0.28 and concordant clinical recommendations^25^ (Fig. 2c). This matters because frontier AI models operate only through cloud APIs not authorized to receive patient data under HIPAA or GDPR and local open-weight alternatives demand heavy GPU infrastructure while sacrificing much of the agentic reasoning that makes these tools useful. Sharing synthetic data (*Z*\*) rather than real data (*Z*) lets a researcher obtain data-aware assistance while the original records never leave the institution.

**Figure 2.**
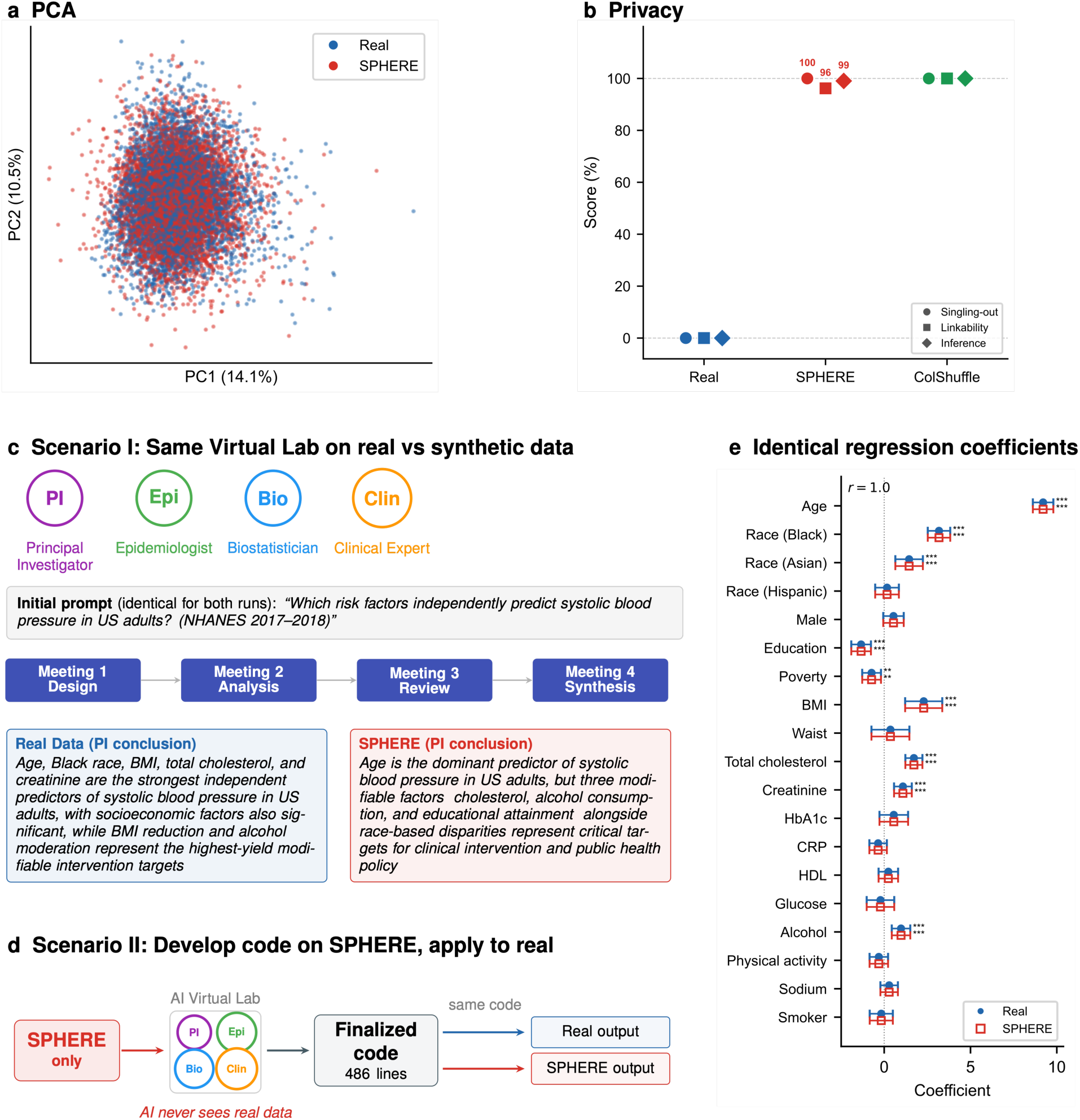
SPHERE enables artificial intelligence (AI)-assisted research without exposing individual-level records. AI Virtual Lab demonstration on NHANES 2017–2018 (n = 4,899; systolic blood pressure). **(a)** Principal component analysis (PCA) of real (blue) and SPHERE (red) records, fully intermixed. **(b)** Anonymeter privacy scores (two-anchor normalized: real = 0%, ColShuffle = 100%); SPHERE reaches near-ceiling privacy on all three metrics (singling-out 100%, linkability 96%, inference 99%). **(c)** Scenario I. Given the identical initial prompt (“Which risk factors independently predict systolic blood pressure in US adults?”), a four-agent virtual lab reaches the same conclusions on real and SPHERE data (R² = 0.28; same top-3 predictors). **(d)** Scenario II. A complete OLS pipeline (486 lines) developed on SPHERE alone, then applied unchanged to both datasets. **(e)** Regression coefficients from that pipeline are numerically identical on real (filled circles) and SPHERE (open squares). Every coefficient matches across all 19 predictors (r = 1.0, maximum difference = 0), and the OLS fit is the same on both datasets (R² = 0.2752).

In Scenario II, where AI never sees the real data, an end-to-end OLS pipeline (486 lines) is developed exclusively on *Z*\* and applied unchanged to both datasets (workflow in Fig. 2d; Supplementary Note 3): *R*^2^=0.2752, coefficient correlation *r* = 1.0, every coefficient, standard error, and *p*-value identical across all 19 predictors (Fig. 2e). The results demonstrate a practical workflow. A researcher shares *Z*\* with any cloud AI service, retains *Z* locally, and verifies returned analyses by running the AI-generated code unchanged on the original data. The workflow requires no modification to existing AI tools, no secure API, and no infrastructure changes at the receiving institution, indicating the privacy protection is entirely on the data side. NHANES 2017– 2018 represents the settings where this workflow matters most, a nationally representative surveillance dataset covering 19 predictors across 12 health domains, with the heterogeneous, multi-source structure typical of clinical and administrative data. The pipeline transfers without modification because SPHERE preserves the same types and statistical structure as the original data, leaving data-cleaning and preprocessing steps unchanged. The complete AI research meeting transcripts for both scenarios (real and SPHERE data runs shown side by side) are provided in Supplementary Note 3.

## Population-scale biobank research is faithfully reproduced with SPHERE

Combining individual-level data across national biobanks would allow analyses to be replicated across populations, extended to larger cohorts and used to train AI models at the scale of millions of participants. The scale, longitudinal depth and genomic, molecular and health-record linkages that make biobanks valuable for AI also make them highly sensitive^26–28^, restricting how they can be accessed, combined and used outside controlled research environments. Access is governed by dataset-specific data-use agreements, ethical approvals and institutional review, and analyses are typically confined to approved research platforms rather than portable computing environments. These platforms support many conventional analyses but limit redistribution, cross-biobank pooling, and the arbitrary large-scale GPU workflows that modern AI development requires. Resources such as UKB, All of Us, FinnGen, and the China Kadoorie Biobank can each support biobank-level discovery but cannot be routinely analyzed together or redistributed for independent reuse. A synthetic release that preserves the analytic behavior of the original data while keeping participant-level records local would overcome this constraint, provided fidelity holds at biobank scale.

We tested SPHERE on two UKB resources: genome-wide genotypes and plasma proteomics (Fig. 3). For the genotype analysis, we retained 9,692,208 bi-allelic imputed variants in 461,307 participants after filtering for INFO ≥ 0.7 and MAF ≥ 0.01. SPHERE was applied within 12 ancestry clusters, jointly transforming AD status, sex and age with the genotype matrix in 2,000-variant blocks (∼75 s per block on a single CPU core; 100 CPU-hours genome-wide). Original and synthetic data were then processed through parallel linkage disequilibrium (LD)-pruning, PCA and genome-wide association study (GWAS) workflows (Fig. S2). LD pruning (MAF ≥ 0.05, r² < 0.1), applied separately to each lane, selected the identical set of 261,793 variants, confirming that SPHERE preserves pairwise LD exactly. SPHERE preserved principal-component (PC) structure to numerical precision, with eigenvalues of the first 40 PCs matched to a maximum absolute deviation of 4.4 × 10⁻^7^, retaining ancestry structure (Fig. 3a, b, c). A GWAS for AD (4,159 cases), adjusting for sex, age and the 40 PCs returned the same effect sizes and significance landscape (Fig. 3d; Supplementary Table 7): per-variant β correlated at r = 1.0 (maximum |Δβ| = 8.4 × 10⁻^9^; maximum |Δ-log_10_P| = 1.8 × 10⁻^6^), genomic inflation was identical (λ = 1.0183; Fig. S3).

**Figure 3.**
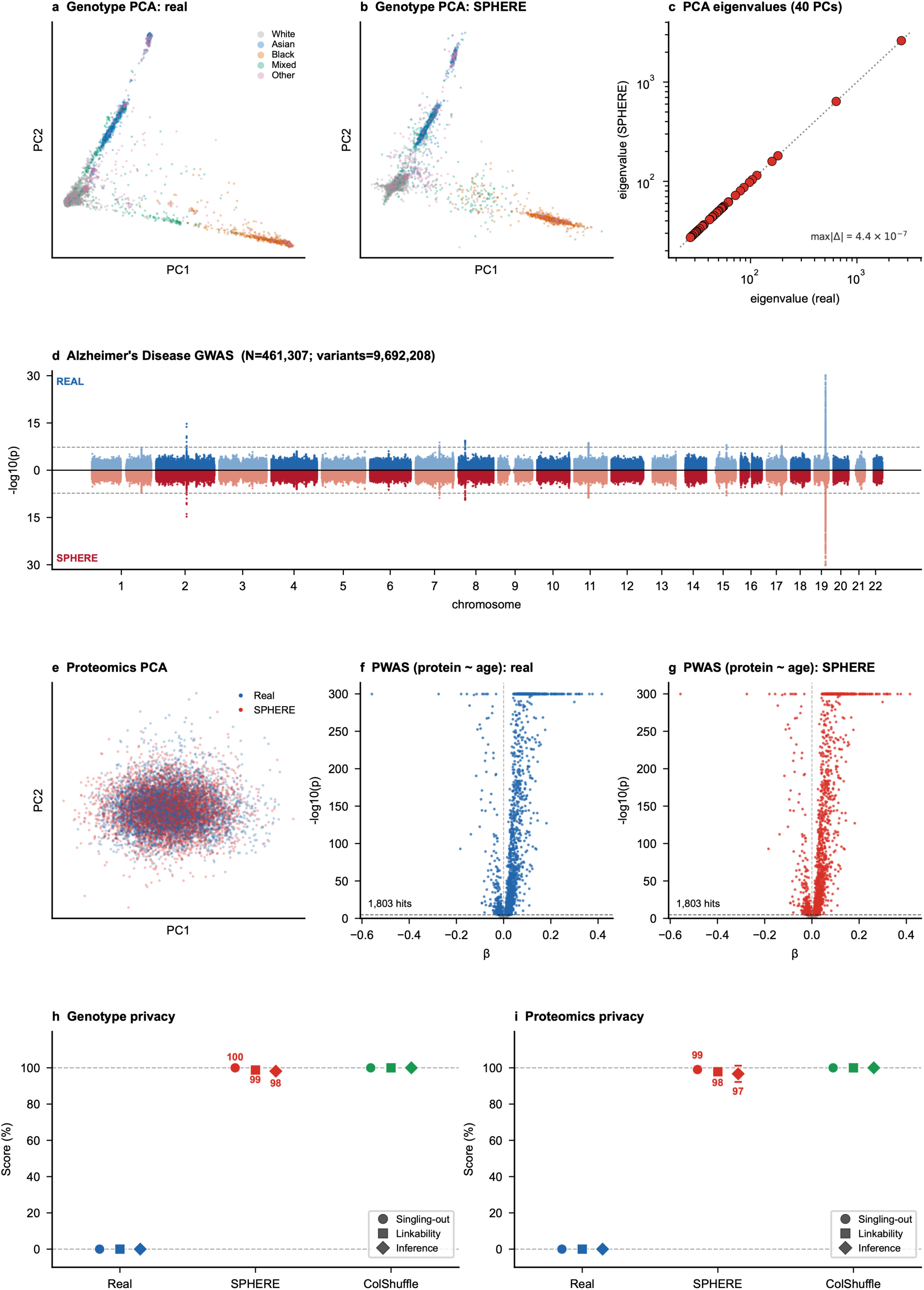
SPHERE reproduces UK Biobank-scale genetic and proteomic analyses without exposing individual-level records. **(a, b)** Principal components of genome-wide genotypes (261,793 linkage-disequilibrium (LD)-pruned autosomal variants, n = 461,307) for (a) real and (b) SPHERE data, colored by genetic ancestry. **(c)** PCA eigenvalues of real versus SPHERE genotypes agree to numerical precision (first 40 PCs; max |Δ| = 4.4×10⁻^7^). **(d)** Miami plot of an Alzheimer’s disease (AD) association study over 9,692,208 imputed variants (4,159 cases) in real (top) and SPHERE (bottom) data; per-variant effects were preserved to numerical precision (β at r = 1.0, max |Δβ| = 8.4×10⁻^9^; −log₁₀P at r = 1.0, max |Δ−log₁₀P | = 1.8×10⁻^6^) with matching genomic inflation (real λ__GC_ = 1.018; sphere λ__GC_ = 1.018) (Fig. S3). −log₁₀P is capped at 30 for display. **(e)** Principal components of the Olink plasma proteome (n = 44,526; 2,916 proteins); real (blue) and SPHERE (red) samples overlap. **(f, g)** Plasma proteome-wide protein–age associations for f real and g SPHERE data, recovering the same 1,803 Bonferroni-significant proteins (P < 0.05/2,916) with matching effect sizes (β at r = 1.0, max |Δβ| < 1×10⁻¹⁵). **(h, i)** Anonymeter empirical privacy scores, scaled from the real-data baseline (0%) to a column-shuffled upper bound (100%) and reported as the mean over 10 draws, for genome-wide genotypes (singling-out 100, linkability 99, inference 98) and the plasma proteome (singling-out 99, linkability 98, inference 97).

Applying SPHERE to the UKB Olink 3k plasma proteome (2,916 proteins), real and synthetic samples were fully intermixed in PC space (Fig. 3e). A proteome-wide association study for age recovered an identical set of 1,803 of 2,916 Bonferroni-significant proteins on real and synthetic data, with per-protein effects matching to numerical precision (β at r = 1.0, maximum |Δβ| < 1×10⁻¹⁵; Fig. 3f, g; Supplementary Table 8). To evaluate machine learning utility, we trained elastic-net model, tree-based ensembles (random forest and gradient boosting) and neural network models to predict age. Elastic-net models retained 100% of the real-data R² and nonlinear alternatives retained 86–95%, most of the predictive utility but without the exact invariance guaranteed for sufficient-statistic analyses (Fig. S4a).

Finally, we quantified empirical privacy risk with the Anonymeter. Synthetic genotypes were strongly protected on all three axes: singling-out 100 ± 0%, linkability 98.8 ± 0.1% and inference 98.2 ± 1.5% (Fig. 3h; Supplementary Table 9); the Olink proteome scored 99.1 ± 2%, 97.8 ± 1% and 96.7 ± 4% for singling-out, linkability and inference (Fig. 3i; Supplementary Table 10). SPHERE thus reproduced population-scale genotype and proteome-wide linear inference to numerical precision while keeping the original participant-level records local and showing low empirical re-identification risk under the evaluated attacks.

## SPHERE enables an open, individual-level release of the Stanford ADRC multi-modal Alzheimer’s disease cohort

Institutions are increasingly conducting deep phenotyping cohorts to integrate neuroimaging, genomics, proteomics, and clinical assessment to enable multi-modal basis for identifying AD subtypes and predicting progression. The United States alone funds more than 30 Alzheimer’s Disease Research Centers (ADRCs), each accumulating multi-modal data on hundreds to thousands of participants. Collectively, they hold data on a scale no single center commands, yet each cohort sits behind institutional review board (IRB) protocols, data-use agreements, and secure enclave requirements that take weeks to months to navigate per site, make cross-center pooling prohibitively slow, and categorically exclude cloud AI tools.

We release a first open individual-level SPHERE version of the Stanford ADRC multi-modal cohort, accessible via portal (https://adrc-sphere.stanford.edu/) that supports browsing, downloading, and autonomous agentic AI analysis under registered access (registration and a click-through data-use agreement), with no IRB review or institutional approval gate. The cohort includes 644 participants across nine modalities: demographics and diagnosis, cognitive assessments, plasma biomarkers, amyloid positron emission tomography (PET), tau PET, SNP genotypes, peripheral blood mononuclear cell (PBMC) single-cell RNA (scRNA), cerebrospinal fluid (CSF) and plasma proteomics. Participants span eight diagnostic groups, including healthy controls (n=290), mild cognitive impairment (n=129), AD (n=79), Parkinson’s disease (n=113; three groups combined), Lewy body disease (n=25) and other diagnoses (n=8). These modalities comprise 206,358 features with 51.9% overall sparsity, reflecting genuine modality-specific missingness rather than uniform measurement across all participants. The SPHERE release preserves this participant-by-modality availability structure (Fig. 4a, Supplementary Note 4). SPHERE preserves exact categorical marginal distributions for all eight diagnostic groups and achieves distributional fidelity ≥ 88% across the eight non-demographic modalities, rising to ≥ 96% for imaging and proteomics modalities (Fig. 4b). The lower KS fidelity for genotype dosage and scRNA reflects the pronounced deviation from the Gaussian distribution. Anonymeter privacy scores, averaged across the eight modalities, are 98.8 ± 1.8% (singling-out), 92.3 ± 4.6% (linkability), and 89.0 ± 10.7% (inference) (Fig. 4c); per-modality scores at *k* = 1 − 5 are given in Supplementary Table 11.

**Figure 4.**
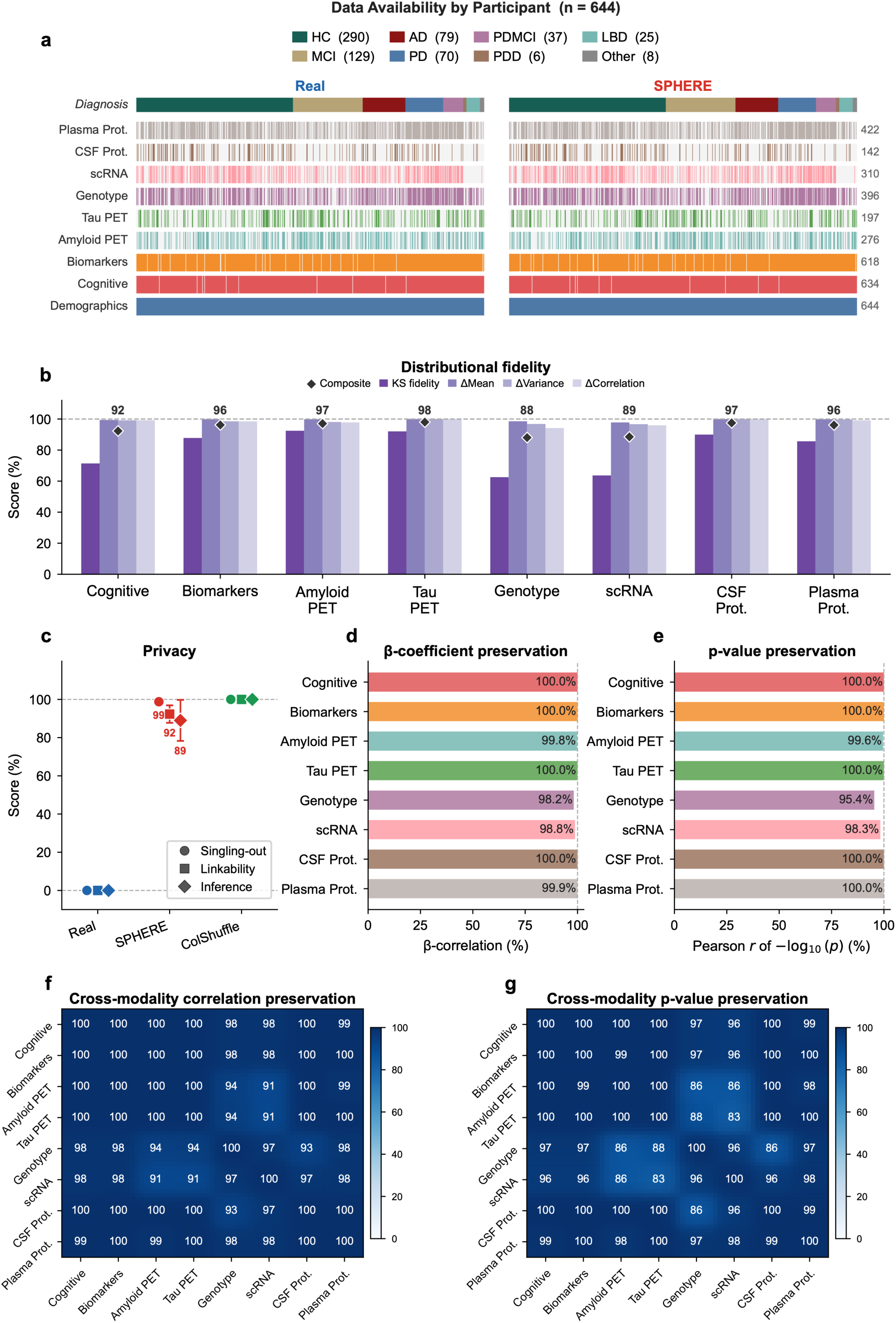
SPHERE reproduces multi-modal analyses on the Stanford ADRC cohort without exposing individual-level records (n = 644; nine modalities). **(a)** Data-availability heatmap, participants sorted by diagnosis; real and SPHERE columns share identical missingness and categorical marginals. (b) Distributional fidelity (per-modality composite of KS statistic and Δmean/Δvariance/Δcorrelation): ≥ 88% for every modality, ≥ 96% for imaging and proteomics. (c) Privacy by two-anchor Anonymeter normalization (real data = 0%, ColShuffle = 100%). SPHERE reaches near-ceiling on all three metrics; singling-out 98.8 ± 1.8, linkability 92.3 ± 4.6, inference 89.0 ± 10.7 (mean ± s.d. across 8 modalities, demographics excluded). (d) OLS β preservation for AD status (50 randomly selected features, 100 replicates): 100% for cognitive, biomarkers, cerebrospinal fluid (CSF) proteomics and tau positron emission tomography (PET); 98.2–99.9% for single-nucleotide polymorphism (SNP) genotypes, scRNA, plasma proteomics and amyloid PET. (e) −log₁₀(p) preservation under the same protocol (95.4–100%, lowest for SNP genotypes). (f) Cross-modality correlation preservation (50 features per modality, 100 replicates): ≥ 90% across all 28 modality pairs, ≥ 97% for same-domain pairs. (g) Cross-modality p-value preservation: 83–88% where SNP genotypes or scRNA pair with amyloid or tau imaging and where SNP genotypes pair with CSF proteomics; all other pairs ≥ 95%, ≥ 99% for same-domain pairs.

Downstream utility is preserved at the level of individual association statistics, despite minor deviations from 100% due to missing data. For OLS regression on AD, the correlation between real and SPHERE *β*-coefficient is 100% for cognitive features, blood biomarkers, tau PET, and CSF proteomics, and 98.2 – 99.9 % for the remaining modalities (Fig. 4d). The corresponding −*log*_10_(*p*) correlations show the similar pattern, reaching 100% for five modalities and 95.4 – 99.6 % for the remaining three with the SNP genotypes lowest (95.4%; Fig. 4e, Supplementary Table 12). Beyond single-modality statistics, SPHERE also preserves cross-modality correlation structure. Averaged over 100 feature subsamples, real versus SPHERE agreement for pairwise cross-modality correlations exceeds 90% across all 28 pairs and reaches 99–100% for closely related domains such as amyloid, tau PET and CSF, plasma proteomics (Fig. 4f). The corresponding significance structure is preserved almost as well. −*log*_10_(*p*) correlations exceed 95% for 23 of the 28 modality pairs and stay above 91% for all but five; those five (83–88%) each involve SNP genotypes or scRNA (Fig. 4g).

Together, these results show that the SPHERE release preserves the within- and cross-modality structure needed for cohort-level analysis. This fidelity makes it possible to expose the cohort not only as downloadable data, but also as an interactive analysis layer. Building on this preserved analysis layer, we implemented an interactive agentic AI portal directly on the SPHERE data. Users can query the cohort and run analyses through natural language, while the agent operates only on the synthetic twin and never receives the original participant records. This separation is essential for agentic workflows, where rule-based restrictions alone can be vulnerable to prompt injection^29,30^, guardrail bypass ^31,32^ and inadvertent disclosure^33^. By grounding the agent in SPHERE data, access to the original records is blocked at the data layer rather than controlled only by policy.

## Landmark studies reproduce end-to-end on SPHERE data

Given SPHERE’s preservation of population structure, association signals and privacy at scale, we examined the reproducibility of scientific conclusions built from the original data. We tested SPHERE at the level of scientific workflows, re-running published analyses from access-restricted cohorts in the UKB, the Stanford ADRC and the Global Neurodegeneration Proteomics Consortium (GNPC)^34^. SPHERE preserved structure rich enough to regenerate not just the linear signals it is designed to carry, but the non-linear clinical endpoints that constitute the actual discoveries.

First, Oh et al.^35^, published in *Nature Medicine*, identified organ-specific aging signatures across the plasma proteome and linked accelerated organ aging to disease and mortality in the UKB. We trained the same organ-aging clock models on SPHERE-transformed proteomics (n = 44,526) and tested whether the resulting organ age gaps reproduced the disease and mortality associations reported from the original data. The conventional clock (trained on the full plasma proteome) reached an identical chronological versus predicted age correlation on real and SPHERE data (r = 0.914 for both; Δr = −1×10^−15^ Fig. 5a), and this held across all 13 organ clocks (maximum |Δr| = 7×10^−15^; Fig. 5b; Supplementary Table 13). The resulting per-organ age gaps carried the same clinical signal. In Cox survival models, each organ’s age gap predicted its target disease with concordant hazard ratios across ten organ–disease pairs (r = 0.967) with concordant estimates for accelerated brain aging and AD (HR = 1.81 real, 1.50 SPHERE), heart aging and heart failure (HR = 1.80 real, 1.54 SPHERE) and lung aging and emphysema/COPD (HR = 1.40 real, 1.26 SPHERE) (Fig. 5c; Supplementary Table 14) and a multi-organ aging burden reproduced a monotone mortality dose–response, from accelerated agers (≥ 8 aged organs, HR = 7.33 real / 5.99 SPHERE) to individuals with youthful organs (HR = 0.58 real / 0.56 SPHERE) (Fig. 5d; Supplementary Table 15). These non-linear endpoints fall outside SPHERE’s exact-preservation class, yet the SPHERE estimates show only modest attenuation toward the null. The direction, statistical significance and rank ordering of effects are preserved.

**Figure 5.**
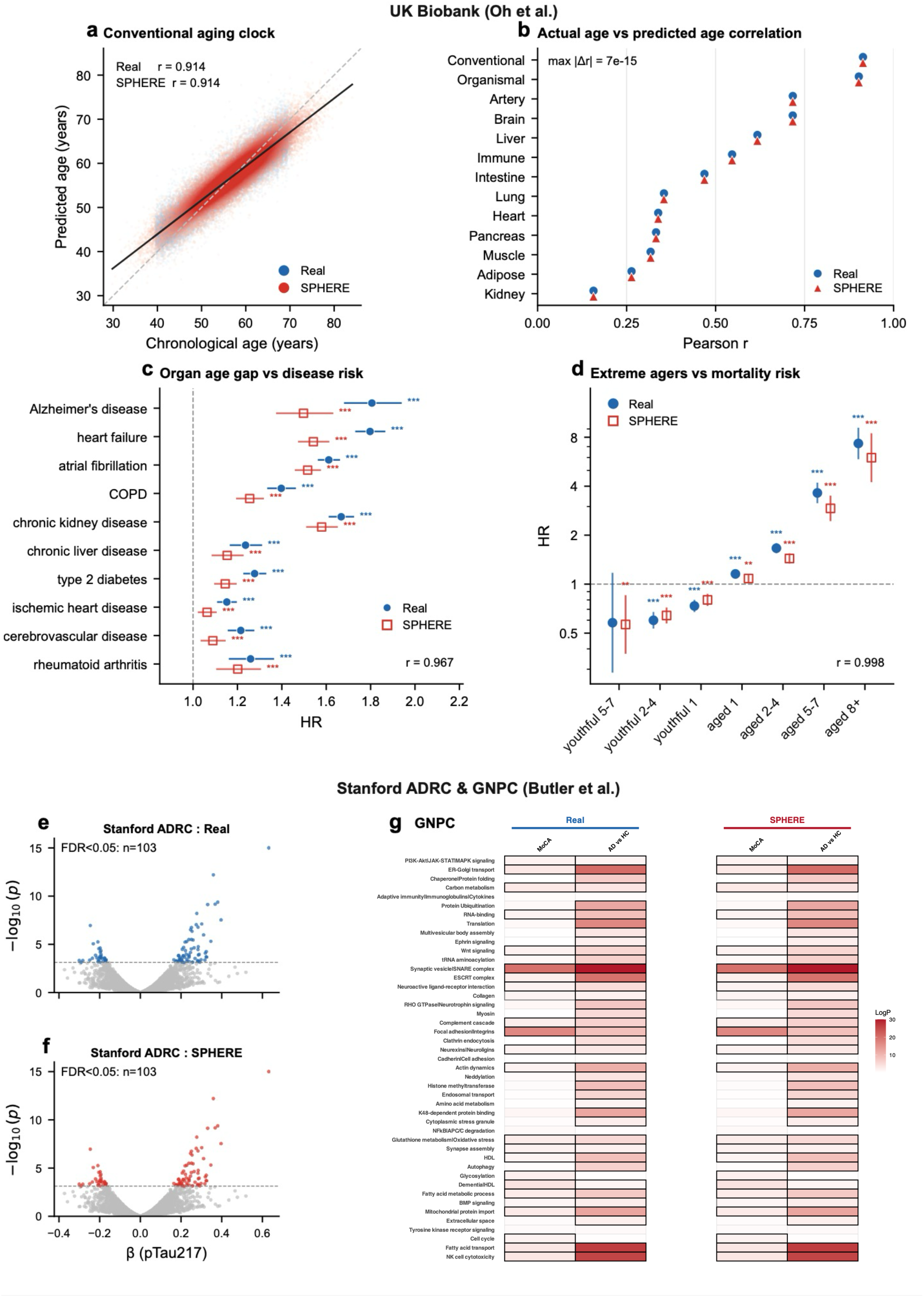
SPHERE reproduces published aging-biomarker analyses across independent cohorts. a–d, UK Biobank (UKB) organ-aging clocks, Oh et al.; e–g, Stanford ADRC pTau217 and Global Neurodegeneration Proteomics Consortium (GNPC) pathway analyses, Butler et al. **(a)** Conventional aging clock (age predicted from the full plasma proteome): predicted versus chronological age on held-out test data; real and SPHERE give a identical fit (r = 0.914 each; Δr = −1×10⁻¹⁵). **(b)** Organ-specific clocks reproduced to numerical precision: test-set Pearson r of predicted versus actual age for 13 organ clocks, real versus SPHERE (max |Δr| = 7×10⁻¹⁵). **(c)** Organ age-gap to incident-disease Cox hazard ratios across 10 organ–disease pairs; log-HR concordance r = 0.967. Wald p (*p<0.05, **p<0.01, ***p<0.001). **(d)** Multi-organ aging burden to all-cause mortality, stratified by count of youthful and aged organs; HR on a log axis, log-HR concordance r = 0.998. **(e,f)** ADRC plasma pTau217 association volcano: real and SPHERE recover the same 103 BH-FDR<0.05 hits with numerically identical effect sizes (β r = 1.0). Dashed line, FDR<0.05 threshold. **(g)** SPHERE data reproduced module score associations with clinical AD diagnosis (AD or HC) or Montreal cognitive assessment (MoCA) in the GNPC cohort. Black boxes indicate significant correlations with significance after FDR adjustment (padj < 0.05).

Second, in an independent cohort and assay, we reproduced the Stanford ADRC plasma proteomic study of Butler et al.^36^, regressing core AD blood biomarkers like phosphorylated tau 217 (pTau217) to the plasma proteome in the Stanford ADRC to identify plasma proteins and functional modules that track with AD progression. These functional modules scores were associated with clinical diagnosis and cognitive decline in the GNPC harmonized dataset. In both datasets, SPHERE recovered exact linear model association, with numerically identical effect sizes. The pTau217 biomarker to plasma proteome linear model retained every one of the 103 proteins significant at FDR < 0.05 (β Pearson r = 1.0; Fig. 5e, f; Supplementary Table 16). In the GNPC, the module-score associations with AD diagnosis (recoded as ±1) and Montreal Cognitive Assessment (MoCA) score reproduced the same clinical signal on SPHERE data (Fig. 5g; Supplementary Table 17). A researcher restricted to SPHERE data reaches the same conclusions as one working from the original records, showing that a SPHERE release carries not only preserved statistical signals but the scientific end products those signals support.

## SPHERE generalizes to deep-learning embeddings across language, vision, and biosignals

Sensitive data blocks not only statistical analysis but AI model training itself. Institutions that hold protected health records, clinical images, or time series recordings can train specialist models locally, but cannot freely share the resulting embeddings or feature representations for collaborative fine-tuning, federated model development, or reproducibility verification. Sharing embeddings is often assumed to be safe, but embedding inversion attacks can recover input tokens from sentence embeddings^37^ and extract demographic attributes from de-identified clinical note embeddings^38^, so shared representations carry a meaningful privacy risk^39,40^. No current approach provides both formal reconstruction privacy and near-lossless downstream model utility for shared neural-network representations, limiting the cross-institutional data diversity that large foundation models require. SPHERE resolves this by applying the same method to the embedding or feature matrix. It preserves all pairwise dot products and optimal linear weights, while giving each row an uncountably infinite set of indistinguishable candidate originals.

We tested SPHERE across three modalities, each representing a distinct foundational architecture family of contemporary AI: Transformer-based sentence encoders for language^41^, convolutional networks (ResNet^42^) for vision, and temporal deep learning models for time series. These three families represent the dominant architectural paradigms of contemporary AI, spanning language encoders, vision encoders, and clinical signal classifiers. Training on SPHERE embeddings reproduces near-identical downstream performance across all three modalities (Table 1; full protocols in Methods and Supplementary Note 5). For language, a GTR (Generalizable T5-based dense Retriever) linear probe on Dreaddit^43^ stress classification reached an area under the receiver operating characteristic curve (AUROC) of 0.795 on both real and SPHERE embeddings (n=715). A ResNet-50 linear probe on ImageNet-100^44^ gave Top-1 accuracy of 77.2±0.07% (real) versus 77.5±0.06% (SPHERE) (n=2,500). The same pattern held for non-linear MLP probes: Dreaddit AUROC held at 0.782 (real) versus 0.780 (SPHERE), and ImageNet-100 Top-1 went from 82.2±0.07% (real) to 80.4±0.08% (SPHERE). Electrocardiogram (ECG) classification on PTB diagnostic ECG database gave 0.727±0.03 (real) versus 0.667±0.03 (SPHERE), where gap reflects the small (∼60-record) PTB test cohort. Downstream performance was similarly maintained across k = 1–5 sequential applications (Supplementary Table 18). We next quantified privacy with Anonymeter (mean across 10 independent evaluation runs). For the text embedding, 98.2% (singling-out 99, linkability 98 and inference 98); for the image embedding, 92.0% (100, 88 and 88) and for the raw ECG waveform, 86.7% (82, 83 and 95). Privacy increased with the number of applications across the full sweep (k=1-5) in all three modalities (Supplementary Table 19). SPHERE preprocessing for all three tasks requires at most ∼17s on a laptop CPU (Apple M3), so the privacy gain carries no substantial computational overhead. Any institution currently sharing embeddings for collaborative AI development, a workflow widely assumed safe despite documented inversion vulnerabilities, can substitute SPHERE representations without modifying any downstream training or evaluation code.

**Table 1.** SPHERE deep-learning experiments across three modalities. SPHERE (k=2) is applied to the embedding or feature matrix before training, and evaluation is always on held-out real data. Values are 10-seed averages ± standard error of the mean (SEM). Privacy is measured by the mean of three re-identification risks (singling-out, linkability, inference), each rescaled on a two-anchor axis where 0% is the real data and 100% a column-shuffled floor averaged over 10 seeds. SPHERE preprocessing time on an Apple M3 CPU (seed 0): text 18ms (2,413 × 769), image 17s (129,395 × 2,148), electrocardiogram (ECG) 17 ms (308 × 12,001). Full experimental protocol in Supplementary Note 5.

| Modality | Dataset | Eval. set | Metric | Real (±SEM) | SPHERE (±SEM) | Privacy |
| --- | --- | --- | --- | --- | --- | --- |
| Text (linear probe) | Dreaddit | test (n=715) | AUROC | $0.795 \pm 0.00$ | $0.795 \pm 0.00$ | 98.2% |
| Text (MLP) | Dreaddit | test (n=715) | AUROC | $0.782 \pm 0.00$ | $0.780 \pm 0.00$ | |
| Image (linear probe) | ImageNet-100 | test (n=2,500) | Top-1 % | $77.2 \pm 0.07$ | $77.5 \pm 0.06$ | 92.0% |
| Image (MLP) | ImageNet-100 | test (n=2,500) | Top-1 % | $82.2 \pm 0.07$ | $80.4 \pm 0.08$ | |
| ECG (Transformer) | PTB | test (n≈64) | AUROC | $0.727 \pm 0.03$ | $0.667 \pm 0.03$ | 86.7% |

## The SPHERE desktop application provides a complete platform for generating, evaluating, certifying, and sharing SPHERE data, with an agentic AI for autonomous research

Each of the five workflows demonstrated above, from AI-assisted research (Fig. 2) and biobank-scale replication (Fig. 3), open cohort release (Fig. 4), end-to-end reproduction of published studies (Fig. 5) to utility-preserving deep-learning training (Table 1), begins with the same steps: generate a SPHERE synthetic twin, verify its privacy and fidelity, and share it. We implement this as a macOS desktop application, available at https://github.com/statzihuai/SPHERE, integrating four modules in a single interface. The Generate module accepts any CSV file by drag-and-drop and produces a SPHERE synthetic twin in seconds using validated default parameters. The Evaluate, Certify and Share module runs the Anonymeter privacy benchmark and distributional fidelity tests, then issues a SPHERE Certificate. The certificate is a cryptographically fingerprinted HTML document embedding the dataset SHA-256 hash, privacy scores, and fidelity scores, and it can accompany a data release or publication as a formal, machine-readable audit record. The SPHERE World module allows users to share and manage SPHERE datasets. Data are stored in user-controlled external storage, and the application never transmits original records. The SPHERE AI module is a locally running agentic AI powered by frontier language model API that operates exclusively on the synthetic CSV inside an OS-enforced macOS Seatbelt sandbox, with no access to the original data by design. Users can confirm reproducibility at any point by running the agent’s accumulated analysis script against their local real dataset, following the produce-on-synthetic, verify-on-real workflow demonstrated in the virtual lab experiment, without transmitting sensitive records to any external service.

## Discussion

We introduce SPHERE, a model-free synthetic-data generator that is the only method among ten evaluated to combine principled privacy, exact preservation of first- and second-order statistics, exact linear inference, competitive machine-learning utility and practical speed. Rather than controlling or perturbing each downstream analysis, SPHERE transforms the data layer itself, producing a representation that can be shared, audited, analyzed by AI agents and used for model development while preserving the analytical structure of the original records. Across five workflows spanning AI-assisted analysis, population-scale biobanks, open cohort release, deep-learning representations and end-to-end reproduction of published biomedical analyses, SPHERE enabled uses of sensitive data that conventional access arrangements make difficult or impossible. Most notably, researchers restricted to SPHERE data recovered not only the original statistical signals but the same biological and clinical conclusions. These capabilities require no model fitting, privacy budget or continuing computational infrastructure.

Beyond the algorithm, we release SPHERE as a desktop application that fully integrates into the study team’s workflow without the need to explicitly code. Four modules span generation of a synthetic twin from any tabular file, evaluation of its privacy and fidelity, issuance of a certificate, and local sharing. The certificate is a cryptographically fingerprinted record embedding the dataset hash together with its privacy and fidelity scores, so any release can carry a machine-readable audit trail that a downstream user can verify independently. An integrated agentic module runs a user-supplied frontier language model on the synthetic data inside an operating-system sandbox that has no access to the original records by design. Its accumulated analysis script can then be re-run on the local real data to confirm reproducibility, the same produce-on-SPHERE, verify-on-real pattern used in the virtual lab. The Stanford ADRC portal is built on this design, offering natural-language analysis of a sensitive cohort directly in the browser, a deployment that would be unsafe on the real records but is sound on their SPHERE twin.

Biomedical research is not unique in adhering to regulations around data. Other sectors like the government, education, and financial --also strive to adhere to regulatory principles. Training AI models on such data requires diversity at a scale that no single institution holds, yet combining data across sites has remained practically infeasible due to regulatory and infrastructural barriers^45–47^. Federated learning partially addresses this by keeping raw data local, but requires coordinated infrastructure at every site, restricts analyses to gradient-based objectives, remains vulnerable to gradient-inversion attacks that can reconstruct training examples^48^, produces no shareable dataset, and cannot flexibly accommodate modified training plans. SPHERE takes a complementary approach: each institution generates a SPHERE file locally and releases it once, with no ongoing infrastructure, no coordination protocol, and a principled privacy guarantee independent of the downstream analysis. Each addition enables cross-population comparisons with every existing corpus, supporting analyses that current data-access frameworks make prohibitively difficult, without requiring bilateral data-sharing agreements (though governance review at each releasing institution remains advisable). The same logic applies outside biomedicine to any domain where institutions hold sensitive tabular data of broad scientific value.

Several limitations define the current scope of SPHERE. First, the exact mathematical guarantee applies only to analyses determined by means, variances and covariances, including ordinary least squares, ridge, lasso, elastic net, principal component analysis, canonical correlation analysis and linear discriminant analysis. Generalized linear models, survival models and tree-based methods fall outside this exact class, although they retained strong empirical performance throughout our benchmark and application studies. Analyses requiring moments beyond second order, including interaction terms and subgroup-specific effects, likewise fall outside the exact class. These attenuate empirically toward the null rather than generating spurious effects, so conclusions drawn from SPHERE data are conservative. Where the data scientist retains the original records this affects only which models are explored, since the resulting analysis code is executed on the real data and returns exact results; where only a SPHERE release is available, the absence of a higher-order effect should not be read as evidence of absence. Second, SPHERE assumes independent observations and therefore does not directly extend to longitudinal repeated measures or graph-structured data with explicit inter-sample dependencies. Third, the formal reconstruction guarantee is stated for an adversary who observes only the released twin. Adversaries holding partial knowledge of original records are addressed empirically rather than formally: the linkability and inference evaluations reported in previous sections assume exactly such an adversary, and SPHERE retains high privacy scores under both. Extending the formal guarantees to these adversary models, and to broader classes of analysis, is an important direction for future work.

SPHERE reframes privacy preservation from protecting individual analyses to protecting the data layer itself. Rather than restricting every downstream computation, a single transformation produces a synthetic representation that can support collaborative analysis, reproducibility, training of AI models and deployment of AI agents while the original records remain protected. As AI becomes increasingly central to scientific discovery, approaches that make sensitive datasets openly usable without exposing participant-level records may help bridge the longstanding divide between privacy and scientific progress.

## Methods

### SPHERE

SPHERE takes a sensitive data matrix with n rows and p columns and returns a synthetic matrix of the same dimensions, in which no row corresponds to any single original participant. It adds no noise, fits no model to the data, and requires no dataset-specific calibration or tuning. A single unified scheme applies to all column types; nominal categorical variables pass through an encode– decode step that reproduces their category counts exactly. Generation time and storage both scale linearly in the number of records, which is what makes the method tractable at biobank scale.

The column means, variances and covariances of the synthetic matrix are identical to those of the original, so any analysis determined by these quantities — including ordinary least squares (OLS), ridge regression, lasso, elastic net, principal component analysis (PCA), canonical correlation analysis and linear discriminant analysis — returns numerically identical results on the two matrices. SPHERE additionally preserves the local per-sample structure that non-linear machine-learning models depend on. The default settings were selected by a sweep across all 33 benchmark datasets (Fig. S1) as the configuration lying on the privacy–utility Pareto frontier. Applying SPHERE k times in sequence (SPHERE×k) strengthens privacy monotonically while leaving the exactness guarantees unchanged, at a measured cost in non-linear machine-learning utility. The algorithmic construction of SPHERE, and the formal derivations underlying the guarantees stated below, are withheld from this preprint pending a patent application (see Code availability).

### Principled privacy guarantee

SPHERE is governed by parameters drawn at random for each synthesis. They are never released alongside the data, are generated afresh for each synthesis and are not retained, so no key material persists after generation.

Without knowledge of these parameters, each released row corresponds to an uncountably infinite possibility of mapping original records. The ambiguity compounds multiplicatively when SPHERE is applied repeatedly: for n = 44,526 and k = 2 (as in the UK Biobank proteomics analysis), the reconstruction set contains at least 7.9 × 10⁹ discrete branches, each spanning an uncountable set of values. SPHERE’s guarantee is therefore an information-theoretic statement about reconstruction ambiguity that holds unconditionally. It operates under a different framework from differential privacy (DP): DP adds calibrated noise scaled to a user-specified budget *ε* (with *δ* the probability the bound fails) to limit information leakage from any computation on the data, whether a query, model training, or synthetic generation; smaller *ε* yields stronger privacy at the cost of greater utility loss. SPHERE is designed for a different setting: sharing a full row-level dataset for downstream analysis. Its appropriate standard is reconstruction ambiguity: the number of original records consistent with each synthetic row. The two frameworks can be combined, but at the cost of exact statistical utility: because SPHERE preserves means, variances and covariances exactly, these aggregates remain recoverable from the released data, violating *ε*, *δ*-DP for any *δ* < 1; satisfying DP requires adding noise that breaks this invariant and with it the exact OLS and moment guarantees. Crucially, the aggregate structure that SPHERE preserves, the first two moments (means and covariances), is the same class of summary statistic routinely shared in practice; in genetics, for example, allele frequencies and linkage-disequilibrium structure are openly released as GWAS summary statistics. SPHERE’s protection is therefore at the level of the individual record, which cannot be reconstructed, and it exposes no aggregate information beyond these conventionally shared summaries.

### Benchmarking

SPHERE, evaluated both at its default single-application setting and as SPHERE×2 (two sequential applications, k=2), was benchmarked against nine comparison methods on 33 real-world datasets from four OpenML benchmark suites (CC18, CTR23, Grinsztajn-cls, Grinsztajn-reg)^49–51^ spanning five scientific domains: life and health sciences (7 datasets), physical sciences and engineering (9), computer science (9), social and behavioral science (4), and food and sensory science (4). Sample sizes range from n=522 to n=10,000 (median 2,109 [908–6,497]) and feature counts from p=5 to p=490 (median 14 [8–25]). The nine methods span the main algorithmic families: ColShuffle (column-shuffle; column-wise independent permutation, privacy and speed baseline); ROMM (dense *n* × *n* orthogonal rotation, a classical microdata-masking baseline); DP-Gaussian (DP, Gaussian mechanism, as deployed in the US Census Bureau 2020 Decennial Census^52^); MVN (classical parametric Gaussian baseline estimating 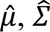, Z^F^ and sampling 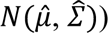; Mixup-*β* (0.4) (row-mixing data augmentation); Synthpop (sequential CART-based synthesis, widely used in official statistics); CTGAN (conditional generative adversarial network); TabDDPM (diffusion model); and GaussianCopula (copula-based synthesis from the SDV library). The last four (Synthpop, CTGAN, TabDDPM, and GaussianCopula) fit a generative model to each dataset; CTGAN and TabDDPM are deep generative models that additionally require hyperparameter tuning and substantial compute and can overfit on small samples. To keep the benchmark tractable across all datasets, every method was run with fixed hyperparameters and no per-dataset tuning. The benchmark therefore measures out-of-the-box rather than peak achievable performance. To bound compute, methods that scale poorly with sample size were not run on the largest datasets: ROMM, CTGAN, TabDDPM, and GaussianCopula were restricted to n≤2,000 (the 16 of 33 datasets), and composite scores are computed over each method’s available datasets. Full parameter settings are in Supplementary Note 2, and per-dataset results are in Supplementary Tables 2-6.

### Evaluation metrics

Each method is evaluated against two references on every dataset. Real data serves as the utility ceiling: a method that matches Real on a utility metric is considered to achieve exact fidelity for that metric. ColShuffle serves as the privacy ceiling and speed reference: it maximally disrupts inter-variable associations while preserving all marginal distributions, and its wall-clock time defines the 1 × speed baseline. Privacy scores are normalized so that ColShuffle = 100 and Real data ≈ 0 (self-disclosure baseline, near-perfect singling-out risk).

**Privacy** is evaluated by adversarial re-identification attacks, following the approach established in the synthetic-data privacy literature^53,54^ and is implemented with Anonymeter (v1.0.0, Python 3.11), which estimates three attack-based risks, each measuring a distinct adversarial capability.

• *Singling-out* (*r_S_*_0_): can an attacker isolate a single individual using a query built from the synthetic data? The attacker picks 3 attribute values from a synthetic row and counts unique matches in the real dataset; success means a real person has been uniquely identified. Risk is the success rate over 500 independent attack attempts.

• *Linkability* (*r_LK_*): if an attacker knows half of an individual’s attributes (set *A*), can the synthetic data reveal the other half (set *B*)? Up to 20 features are selected at random and split evenly into A and B, and the halves are linked by nearest-neighbor matching with n_neighbors = 1 (the Anonymeter default); a success is a target whose nearest neighbor under A and under B is the same synthetic record. Risk is the success rate over 500 independent attack attempts.

• *Inference* (*r_INF_*): can an attacker who knows all of an individual’s attributes except one (the “secret”) predict that secret using the synthetic data as a training set? A single randomly chosen column serves as the secret; risk is the success rate over 500 independent attack attempts.

Each risk is baseline-adjusted (the attacker’s success in excess of a random-guessing control), clipped to [0,1] with 0 denoting no risk above random and 1 fully exposed. Each risk is normalized separately with Real data (≈ 0, worst) and ColShuffle (100, best) as per-dataset anchors:

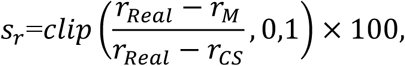

where *r_M_* is the risk of the evaluated method, and *r_Real_* and *r_CS_* are the corresponding risks of the real data (worst-privacy anchor) and ColShuffle (best-privacy anchor), respectively. The composite Privacy score is the mean of the three separately normalized values. To make the stochastic attacks reproducible, all internal random-number states were re-seeded before each evaluator run.

**Distributional fidelity** is quantified by four statistics that compare each column of the synthetic matrix *Z*^∗^ with the corresponding column of the real matrix *Z*. Three are percentage deviations of the column-wise moments, the mean 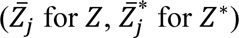, the variance 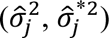, and the *p* × *p* sample correlation matrix 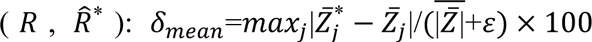, 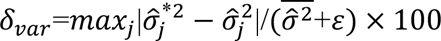, and 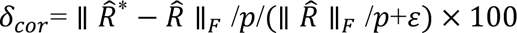, where *j* = 1, …, *p* is the column index, *ε*=10^−8^ and 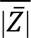 and 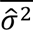 are the mean of 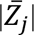 and of 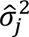 over all columns of all 33 datasets, respectively. The fourth is the mean column-wise Kolmogorov–Smirnov (KS) statistic 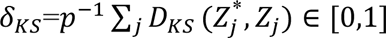, scored as (1 − *δ_KS_*) × 100%. Each of the three percentage statistics is scored as *clip*(100 − *δ*, 0,100), so Real data (*δ* = 0) scores 100 and a method loses one point per one percentage point of deviation.

**Statistical inference utility** is evaluated by fitting OLS to simulated null (*β* = 0) and signal (*β* ≠ 0) responses on each synthetic dataset, averaged over five replicates per dataset. Three outcomes are computed from the coefficient t-statistics: Type I error 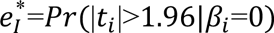, power *pow*\*=*Pr*(|*t_i_*|>1.96|*β_i_* ≠ 0), and 95% confidence interval (CI) coverage 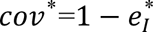. Each is scored relative to Real data’s observed value on the same dataset (not against a nominal target). For higher-is-better metrics (power, CI coverage): *score*=*clip*(1 − *max*(0, *R* − *S*)⁄|*R*|+*ε*, 0,1) × 100, giving 100 when *S* ≥ *R* and falling proportionally as *S* drops below *R* (e.g. *S* = *R*/2 ⇒ 50), where *R* is Real data’s observed value of the metric on that dataset and *S* is the synthetic method’s value. For the lower-is-better metric (Type I error): *score*=*clip*(1 − *max*(0, *S* − *R*)⁄|*R*|+*ε*, 0,1) × 100, giving 100 when *S* ≤ *R* and 0 when *S* ≥ 2*R*.

**Machine-learning (ML) utility** uses a Train-on-Synthetic, Test-on-Real (TSTR) protocol: five regression models were trained on the synthetic dataset and evaluated on a held-out real test set (Elastic Net (EN; α = 0.01), Support Vector Machine (SVM; radial-basis-function kernel), Random Forest (RF; 50 trees), Gradient Boosting (GB; 50 stages) and a Multi-Layer Perceptron (MLP; two hidden layers of 64 and 32 units, max_iter = 1000, early_stopping = True; n_iter_no_change = 20). Elastic Net was given max_iter = 2000 to ensure convergence; all remaining hyperparameters are scikit-learn (v.1.9.0) defaults, and RF, GB and MLP were fitted with a fixed random seed. EN, SVM and MLP were trained on the standardized target and their predictions mapped back to the original scale before scoring. RF and GB were trained on the untransformed target. For each model-dataset pair, score 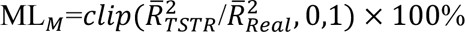, where each 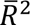 is averaged over five random 8:2 splits; pairs with 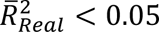 are excluded. Each dataset’s ML utility is the mean score across its remaining models. Of the 33 candidate datasets, one (yprop_4_1) fell below the threshold for all five models and was excluded entirely; per-model exclusions ranged from 1 (RF, GB) to 6 (SVM) and correspond to datasets on which even the real-data model had no predictive signal (Supplementary Table 5).

**Computing time** is reported as, the mean wall-clock time relative to ColShuffle. For each method *M*, we measured the wall-clock time *T_M_* to generate one synthetic dataset on a single CPU core (Apple M3) and report it relative to the column-shuffle baseline as the ratio *T_M_*/*T_ColS_*_ℎ*uffle*_, averaged across the 33 datasets (ColShuffle ≡ 1×).

### NHANES Virtual Lab study

The NHANES 2017–2018 dataset was obtained from the CDC public repository. The analytic sample comprised 4,899 adults with complete observations across 19 predictors (13 continuous and 6 binary) from 12 survey components. SPHERE was applied at its default settings (a single application) to the combined *n* × 20 feature–outcome matrix. The AI Virtual Lab consisted of four specialist agents (a Principal Investigator, Epidemiologist, Biostatistician, and Clinical Expert) powered by Claude Sonnet 4.6, accessed through the Anthropic API. The agents worked through a four-meeting protocol (Study Design, Statistical Analysis, Results Review, Synthesis). In Scenario I, both runs (real and SPHERE) used identical initial prompts. In Scenario II, the lab developed its analysis code exclusively on *Z*\* and the resulting 486-line pipeline was applied unchanged to both *Z*\* and *Z*, so the agents never accessed real data at any stage. Both scenarios’ full transcripts are provided in Supplementary Note 3.

### UK Biobank population-scale study

Genotype data were obtained from the UK Biobank imputed autosomal dataset (Category 100319), comprising 461,307 participants. For the Alzheimer’s disease (AD) genome-wide association study (GWAS), variants were filtered to imputation INFO ≥ 0.7 and minor allele frequency (MAF) ≥ 0.01, yielding 9,692,208 variants. For genotype analyses, SPHERE was applied within ancestry clusters to prevent mixing across genetically distinct groups. Twelve clusters were defined from the genetic principal components (Category 22009; PC1-40) using Ward’s minimum-variance hierarchical clustering based on Euclidean distance (SciPy v1.17.1)^55^. The clustering tree was fit on a random subsample of 40,000 participants; remaining participants were assigned to the nearest cluster centroid, and clusters containing fewer than 1,000 participants were merged with the nearest larger cluster. The resulting clusters ranged from 1,815 to 321,739 participants. SPHERE was applied independently within each cluster using two sequential SPHERE applications (k = 2). AD status (Category 24021; 4,159 cases), encoded as −1 for controls and 1 for cases, together with sex and age were transformed jointly with the genotype matrix. For computational scalability, genotype variants were processed in consecutive column blocks and in parallel. Because the same transformation was applied to every block, this procedure is equivalent to transforming the complete genotype matrix at once. For principal component analysis (PCA), the original and SPHERE-transformed genotype matrices were processed independently. In each matrix, variants were restricted to MAF ≥ 0.05 and linkage disequilibrium (LD)-pruned using a streaming NumPy implementation with a 50-variant window, a five-variant step and an r^2^ threshold of 0.1. The first 40 PCs were then estimated independently from the two pruned genotype matrices using materialized randomized singular value decomposition based on Halko et al.^56^. For the AD genome-wide association study (GWAS), the original-data GWAS was adjusted for sex, age and the first forty PCs estimated from the original genotype matrix. The SPHERE-data GWAS used the corresponding transformed sex and age variables and the first 40 PCs estimated independently from the SPHERE genotype matrix. Both datasets were analyzed using the same vectorized ordinary least-squares pipeline implemented in NumPy (v1.26.4), with covariate adjustment by Frisch–Waugh–Lovell projection and two-sided t-test p-values calculated using SciPy (v1.17.1).

For plasma proteomics, analyses were performed using 44,526 UKB participants profiled with the Olink Explore platform (2,916 NPX-normalized proteins). SPHERE (default settings; k = 2) was applied to protein matrix for both PCA and proteome-wide association study (PWAS) comparisons. Original and synthetic datasets were analyzed using an identical downstream workflow. PWAS was performed by regressing each protein on chronological age using vectorized OLS without additional covariates implemented in NumPy and SciPy, with statistical significance determined using Bonferroni correction (0.05/2,916).

For proteomic machine-learning utility across SPHERE iterations, we sampled 8,000 participants and 300 proteins and predicted chronological age from the protein profile. Utility was assessed using a TSTR framework: EN, RF, GB and MLP were first trained on real data to fix a baseline test-set R², then retrained on SPHERE data at each iteration k = 1–5 and rescored on the held-out real test set. We report the mean and standard deviation across five independent 8:2 splits. The MLP comprised two hidden layers with 128 and 64 units, respectively, each followed by batch normalization and a ReLU activation, with dropout of 0.5. It was trained using Adam (learning rate, 0.001; weight decay, 0.01), with 15% of the training data reserved for internal validation, early stopping with a patience of 25 epochs, a maximum of 500 epochs and a batch size of 256 (PyTorch v2.12.1). EN, RF and GB used the same hyperparameter settings as in the benchmark analyses.

Privacy was evaluated for both genotype and plasma proteomics using Anonymeter. For genotype data, privacy evaluation was performed on the 261,793 LD-pruned variants shared between the original and SPHERE datasets. Because SPHERE was applied within ancestry clusters, each of the 12 clusters was evaluated separately across 10 independent draws. In each draw, up to 2,000 participants were sampled per cluster together with an attacker pool of 200 variants; the two clusters containing fewer than 2,000 participants (n = 1,947 and n = 1,815) contributed all available participants. Singling-out attacks used three-variant queries over the 200-variant pool, whereas linkability and inference each used 20 randomly sampled variants, configured as two 10-variant auxiliary sets for linkability and as one secret plus 19 auxiliary variants for inference. For each draw, scores were first averaged across the 12 ancestry clusters, and genotype privacy is reported as the mean ± s.d. across the 10 draw-level means. For plasma proteomics, an equally sized subsample (n = 2,000) and the same Anonymeter attack protocol were applied to the full 2,916-protein matrix without a variant pool. Scores are reported as the mean ± s.d. across the 10 draws.

### Stanford ADRC data portal

Each participant’s most recent visit was retained to produce a single cross-sectional row. Variables not recorded at that visit were filled using the most recent prior non-missing observation for that participant. Categorical variables (sex, race, ethnicity, diagnosis) were encoded as numeric contrast columns (24 columns total). Participants were partitioned into blocks defined by their modality-presence patterns (9-bit binary vectors). Blocks with fewer than 10 participants were merged iteratively using a globally optimal cheapest-pair purity cost strategy, at each step selecting the merge that minimizes cross-modal contamination weighted by modality rarity, until every block held at least 10 participants. Missing entries were temporarily imputed to global column means before each SPHERE application (default settings; *k* = 2 applications) and the original NaN mask was restored afterwards. Because of the missingness, the released per-modality files preserve second-order structure closely but not exactly (Fig. 4d, e). Per-modality CSV files were extracted in their original measurement scale and published at https://adrc-sphere.stanford.edu/, an open portal supporting data browsing, data download, and interactive agentic AI analysis of the synthetic cohort. Privacy was evaluated using Anonymeter with two-anchor normalization (real data as 0%, column-shuffled as 100%). Utility was evaluated via within-modality OLS *β*-correlation and *p*-value preservation (outcome: ±1 AD indicator) and cross-modality correlation preservation across all 28 modality pairs. Both used 50 randomly selected features per modality, averaged over 100 feature-subsample seeds. Full details are in Supplementary Note 4.

### UK Biobank organ aging proteomics study

Plasma proteomics (Olink Explore, NPX-normalized) covering 2,916 QC-passing proteins in 44,526 participants, ICD-defined incident-disease endpoints, and all-cause mortality follow-up were obtained from the UKB. Organ-enriched protein sets, endpoint definitions, and model training followed Oh et al.^35^. We reproduced their organ-aging pipeline. For each of 11 organs, together with a conventional (all-protein) and an organismal (non-enriched) model, we trained a LASSO age predictor (scikit-learn 1.9.0 LassoCV). The organ age gap (Δage_z) was the z-scored residual of predicted age regressed on chronological age. To test reproduction, we re-fit the identical pipeline on SPHERE×2 (k=2) proteomics and evaluated it. Cox proportional-hazards models (lifelines 0.30.3 CoxPHFitter) related each organ age gap to its incident disease, with age as the timescale, left truncation at baseline, prevalent cases excluded, and sex as a covariate. Real and SPHERE estimates were compared by the Pearson correlation of log hazard ratios across organ–disease pairs. Multi-organ burden was the count of accelerated organs (Δage_z ≥ 1.5 s.d.) banded as 1, 2–4, 5–7 and ≥ 8, and symmetrically the count of youthful organs (Δage_z ≤ −1.5 s.d.) banded as 1, 2–4 and 5–7, each against a no-extreme-organ reference.

### Stanford ADRC and GNPC plasma proteomics

Data and analysis followed Butler et al. Stanford ADRC plasma was assayed on the SomaScan 7k panel, and we tested the association between plasma pTau217 and each protein by linear regression in R 4.4.3 on inverse-normal-transformed values (RNOmni 1.0.1.2), adjusting for the demographic, ancestry and technical covariates and the first six per-protein surrogate variables (sva 3.52.0) of Butler et al. The GNPC Harmonized Dataset V1 (AD Workbench), filtered to SomaScan 7k baseline samples, comprised 11,042 participants (5,635 cognitively healthy, 2,704 mild cognitive impairment, 2,703 AD), with MMSE harmonized to MoCA as in Roheger et al.^57^ Functional proteomic modules from Butler et al.^36^. were scored by a weighted sum of scores and tested against AD diagnosis (coded ±1) and cognitive score by the same linear regression, adjusting for age, sex, contributor group and race. In both analyses SPHERE×2 was applied, and actual versus SPHERE associations were compared by the Pearson correlation of the test statistic.

### Deep-learning embedding experiments

In all three experiments, SPHERE×2 (k=2) was applied jointly to the training-set embedding or feature matrix and the corresponding ±1-coded label vector or matrix. Validation and test sets always used the original representations. Downstream utility was evaluated using a TSTR framework, and results are reported as the mean ± standard error of the mean (SEM) across 10 repeated random seeds or data splits.

**Text (Dreaddit stress classification).** The Dreaddit corpus^43^(Reddit posts labelled for binary psychological stress; HuggingFace andreagasparini/dreaddit) provides a 715-post test split. The training split is randomly partitioned, into 2,413 training and 425 validation examples. Sentence embeddings are extracted once with the frozen gtr-t5-base encoder (sentence-transformers, d = 768) and cached. A linear probe and a shallow multilayer perceptron (MLP) were trained on the transformed training embeddings and continuous transformed labels, and evaluated on the original validation and test embeddings using their binary labels. Performance was assessed using test AUROC averaged across ten random seeds.

**Image (ImageNet-100).** ImageNet-100 consists of the 100 lowest-index classes of ImageNet: 129,395 training images and 5,000 validation images (50 per class). A frozen ResNet-50^42^ (IMAGENET1K_V1 weights) extracts 2,048-dimensional features from its final global average-pooling layer; the backbone is frozen, and the features are computed once and cached. Because ImageNet test labels are not public, the 5,000 official validation images are re-split into two class-stratified halves: a 2,500-image validation set during training and a 2,500-image test set for final evaluation. A linear probe and a shallow MLP were trained on the transformed training features and continuous transformed multiclass labels, and evaluated on the original validation and test data. Performance was assessed using Top-1 accuracy averaged across ten random seeds.

**ECG (PTB Diagnostic Database).** The PTB Diagnostic electrocardiogram (ECG) Database ^58,59^ contains 12-lead recordings (448 recordings; 368 myocardial infarctions (MI), 80 healthy controls (HC)) from 200 patients (148 MI; 52 HC). We performed 10 repeated random patient-level splits with a 70/15/15 train/validation/test ratio, stratified by diagnosis. All recordings from a given patient were assigned exclusively to one subset to prevent data leakage. A Temporal Transformer classifier was trained on the transformed training waveforms from scratch and continuous transformed labels, and evaluated on the original held-out recordings using their binary labels. Performance was assessed using test AUROC averaged across ten random seeds.

**Privacy evaluation.** We evaluated the text embeddings, image embeddings and ECG waveforms after SPHERE transformation. Risks were averaged over 10 runs, each drawing 20 features at random, with singling-out evaluated over predicates of three columns and up to 50,000 attempts per attack, and were reported across k=1 to 5 sequential SPHERE applications.

### SPHERE desktop application

The SPHERE macOS desktop application (Electron; TypeScript/React; distributed as a signed ARM64 DMG, app identifier edu.stanford.sphere) implements the full workflow in four integrated modules. The Generate module applies the SPHERE algorithm to an arbitrary CSV, streaming per-row progress and writing the synthetic twin to disk with no user-specified hyperparameters required beyond the validated defaults. The Evaluate, Certify & Share module invokes a compiled Python sidecar that runs the full Anonymeter attack battery (singling-out, linkability, and inference) alongside distributional fidelity tests; on completion it renders a SPHERE Certificate, an HTML document embedding the SHA-256 fingerprint of both the real and synthetic CSVs, all privacy and fidelity scores, and generation parameters, that serves as a portable audit record for data releases or publications. The SPHERE World module allows users to share and manage sharing of SPHERE datasets via user-controlled external storage (S3 or Dropbox); the application writes files directly to user-owned storage, retains no copy, and never relays original data to external servers. The SPHERE AI module launches a multi-turn Claude agent (Anthropic Claude; users supply their own API key) whose data access is strictly limited to the synthetic CSV: it is passed via the SPHERE_CSV environment variable into a macOS Seatbelt sandbox profile that enforces network isolation, restricts filesystem writes to a temporary working directory, and prevents subprocess spawning beyond the sandboxed Python interpreter. The agent accumulates a reproducible analysis.py across conversation turns; deploying this script on the real data locally verifies concordance without transmitting sensitive records. This sandboxed architecture eliminates the prompt-injection and data-exfiltration risks that arise when agentic AI operates directly on sensitive data, providing a structural privacy guarantee independent of safety guardrails or policy restrictions.

## Data availability

All datasets used in the empirical evaluation are publicly available from OpenML (https://www.openml.org). The Dreaddit corpus is publicly available from the HuggingFace Hub (andreagasparini/dreaddit). ImageNet-100 is derived from ImageNet, available under its terms of use at https://www.image-net.org. The PTB Diagnostic ECG Database is publicly available from PhysioNet (https://physionet.org/content/ptbdb/). The NHANES 2017–2018 data are publicly available from the CDC National Center for Health Statistics (https://wwwn.cdc.gov/nchs/nhanes/). UK Biobank data are available upon request to qualified researchers through a standard protocol (https://www.ukbiobank.ac.uk/register-apply). The Stanford ADRC data used in this study are available upon reasonable request to the Stanford ADRC Data Release Committee (https://web.stanford.edu/group/adrc/cgi-bin/web-proj/datareq.php). GNPC data are available upon request to qualified researchers through a standard protocol (www.neuroproteome.org/harmonized-data-set-hds). Stanford ADRC SPHERE synthetic data are available under registered access (registration and a click-through data-use agreement, with no approval process or committee review) via a portal supporting browsing, downloading, and interactive agentic AI analysis at https://adrc-sphere.stanford.edu/; the original clinical data are available to qualified researchers through the Stanford ADRC data access process.

## Code availability

The SPHERE macOS desktop application is available at https://www.sphereworld.ai/; https://github.com/statzihuai/SPHERE. SPHERE is described in this preprint at the level of what it preserves and what it achieves; the algorithmic construction and the formal derivations are withheld pending a patent application. Both are available to editors and referees on request and will be disclosed in full in the peer-reviewed publication.

## Acknowledgements

Stanford ADRC data collection was supported by the Stanford Alzheimer’s Disease Research Center, NIH/NIA grant P30 AG066515. Research on the Alzheimer’s Disease application was supported by NIH/NIA awards AG089509, AG066206, and AG066515 (Z.H.).

## Declaration of AI use

The authors used Claude (Anthropic) to assist with code development, literature search, and manuscript editing. All theoretical results, simulations, and scientific conclusions were developed and verified by the authors.

## Author contributions

Z.H. conceived the method, developed the theory, implemented the code, performed the experiments and wrote the manuscript. J.P. verified the code, improved experiments, and revised the manuscript. R.C.P. and J.L. processed and analyzed Stanford ADRC data. X.Z. provided critical comments on the deep learning section. R.R.B. and A.W. performed analysis of ADRC and GNPC proteomics data. R.R.B., A.W., L.T., J.W., S.S., E.M., T.W.-C., V.W.H., F.M.L., M.D. and R.A. provided critical review of the manuscript.

